# Structural studies on cell wall biogenesis-I: A method to produce high-quality protoplasts from alga *Micrasterias denticulate*, an emerging model in plant biology

**DOI:** 10.1101/2021.06.30.450603

**Authors:** M. Selvaraj

## Abstract

*Micrasterias denticulate* is a freshwater unicellular green alga emerging as a model system in plant cell biology. This is an algae that has been examined in the context of cell wall research from early 1970’s. Protoplast production from such a model system is important for many downstream physiological and cell biological studies. The algae produce intact protoplast in a straight two-step protocol involving 5% mannitol, 2% cellulysin, 4mM calcium chloride under a ‘temperature ramping’ strategy. The process of protoplast induction and behavior of protoplast was examined by light microscopy and reported in this study.

## Introduction

*Micrasterias denticulate* is a flat disc-shaped freshwater unicellular green algae of size near to 200μm diameter. Taxonomically, this alga belongs to order Desmidales (family: Desmidiaceae), under Streptophyta that are close to land plants in terms of evolution. Morphologically, the alga has a bilateral symmetry with two half-cells or semicells connected at the center, one younger than the other. The two semicells are joined at a narrow neck like central constriction termed as ‘isthmus’. Each mature semicell has one polar lobe with four lateral lobes on its left and right side (Figure 1). In its side view, the alga is fusiform. On the flat surface of the cells there are cell pores that secrete mucilage polysaccharide. The nucleus covers isthmus area and a large flat chloroplast with pyrenoid occupies the centre of each semicells. During mitosis that occurs for every 3-4 days once, the cell divides at isthmus and proceeds through defined stages like bulge stage, two-lobe, five-lobe stage, lateral lobe doubling etc that can be identified morphologically under optical microscope (Meindl, 1993; 2016). At ultrastructural level, distinct stages like septum formation, primary wall stage with spherical vesicles approaching plasma membrane, secondary wall stage with flat vesicles near plasma membrane, and a later stage with mucilage filled pore vesicles at cell pores were noticed (Dobberstein and Kiermayer, 1972; Keirmayer, 1970, 1981; Meindl *et al.*, 1993, Staehelin and Kiermayer, 1970).

**Figure 1:**
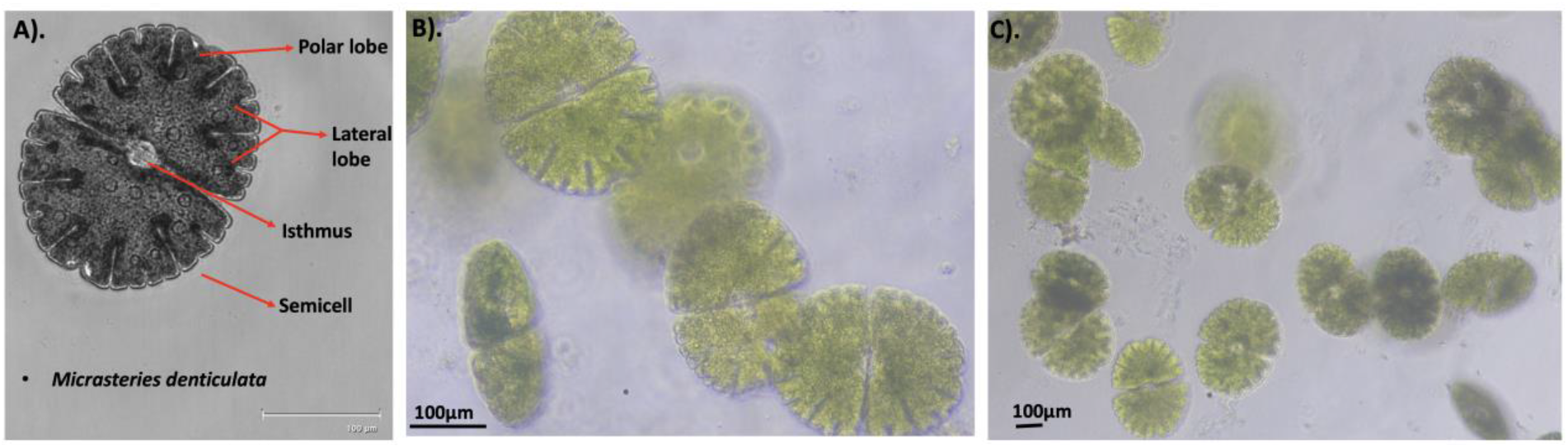
A) Morphology and overall view of *M.denticulata* with its lobes and isthmus marked. (B) and (C) are the status of 3-weeks old culture cells at different magnifications examined before the experiments. One cell showing its fusiform shape in side view is seen in B near scale bar. The scale bar for colour images are drawn based on the scale bar and cell size from the black and white images captured in FLoid cell imaging system along with a scale bar.

Waris, with a defined a culture medium for the cultivation of this algae initiated a series of cytological studies on this desmid (Waris 1950a, 1950b, 1953; Waris and Kallio, 1964). This was followed by Kiermayer, and by Meindl. In the laboratories of Kiermayer, Meindl and others, this algae was used to study critical plant development processes like cell morphogenesis, cell wall biosynthesis, cellulose arrangement, tip growth, metal toxicity, abiotic stress responses etc., establishing this desmid algae as an emerging model in plant cell morphogeneisis and cell biology (Kiermayer, 1964, 1967, 1968, 1970a, 1970b, 1980, 1981; Kiermayer and Meindl, 1980, 1981, 1984, 1986, 1989; Kim *et al.*, 1996; Lacalli, 1975; Meindl and Salomon, 2000; Meindl *et al*, 1990, 1992, 1994, 2016). Since this alga was under investigation from early 1970s, many were interested in producing protoplast from this algal species and unsuccessful attempts to obtain them had been reported (Tippit and Pickett-Heaps, 1974). Subsequently, Berliner and Wenc (1976) developed a protocol for isolation of protoplast from desmids *Cosmarium turpini, Micrasterias denticulata* and two other species *Micrasterias thomasiana* and *M.angulosa* (Berliner and Wenc, 1976).

Berliner and Wenc found that incubating these diverse algal cells with 0.3M mannitol (5.5% w/v solution) and 2% cellulysin at room temperature yielded protoplast. With 12 figures, their experimental outcomes were described in their article about production of protoplasts in all their specimens except *M. denticulate* (Berliner and Wenc, 1976). In the present study, these experiments were repeated with *M.denticulata* with an objective to obtain intact protoplast from this particular species. Such protoplasts are the starting material for a closer examination and following of cell wall biogenesis at high-resolution. It was found that high-quality intact protoplast could be produced reproducibly from *M. denticulate* with a slight necessary modification to Berliner and Wenc protocol, particularly to aid protoplast release, that otherwise locks up inside the cell.

## Materials and Methods

### Culture conditions

Cells of *M.denticulata* were received from the laboratory of Prof.Ursula Lütz-Meindl, University of Salzburg. The cells were inoculated in desmidian medium (supplementary material) with soil extract included and cultivated for 3 weeks with a 14/10-hours light-dark regime at 20 **°**C. The cells were then examined in light microscope to ensure algal growth by looking for its various growth stages and health before protoplast induction experiments. A FLoid cell imaging station and a Nikon Eclipse Ts2 microscope was used for this examination.

### Protoplast induction

Cells from 100ml culture were pelleted by a short spin for at 5000rpm in 50mL falcon tubes. The pellet was suspended in distilled water and vortexed briefly to wash the mucilage on cell surface and pelleted again. The resulting pellet was suspended in 5mL desmidian medium and used for the experiment. Mannitol and cellulysin stocks were made on desmidian medium. Protoplast induction experiments were carried out in 24-well cell culture plates where algal cells, mannitol and cellulysin were mixed and incubated at 22 **°**C. 4mM of calcium chloride was included as many plant protoplast induction experiments use calcium. The cells were examined for 5 hours after each hour and left overnight in darkness undisturbed.

## Results

### *M.denticulata* arrest at lobing, round up stages of protoplast induction at 22 °C

In a search grid to identify a condition that induce protoplast from *M. denticulate*, the cells were mixed with mannitol in a range of concentration from 3% to 9% and cellulysin from 2% to 6% with 4mM calcium chloride included and incubated at 22 **°**C in darkness. After two hours of incubation, most of the cells begins to undergo plasmolysis and starts to retract the protoplast from the cell wall. In 5 hrs time, almost all cells plasmolyzed and their protoplast are highly lobed, irrespective of mannitol concentration (Figure 2A). In conditions with 4 to 9 % mannitol, apart from lobing, in many cells, the protoplast rounds up in each semicell and remain connected at the isthmus as two circular structures touching each other (Figure 2B-C). This round up stage didn’t proceed further to protoplast release, even after overnight incubation. There was no difference in the appearance of cells with variable amount of cellulysin in the inducer medium. Attempts to release the protoplast from these cells by mechanical means like slight sonication or by Dounce homogenizer resulted in cell debris. Continuous shaking of the plate during induction process in the incubator also did not aid for protoplast release.

**Figure 2:**
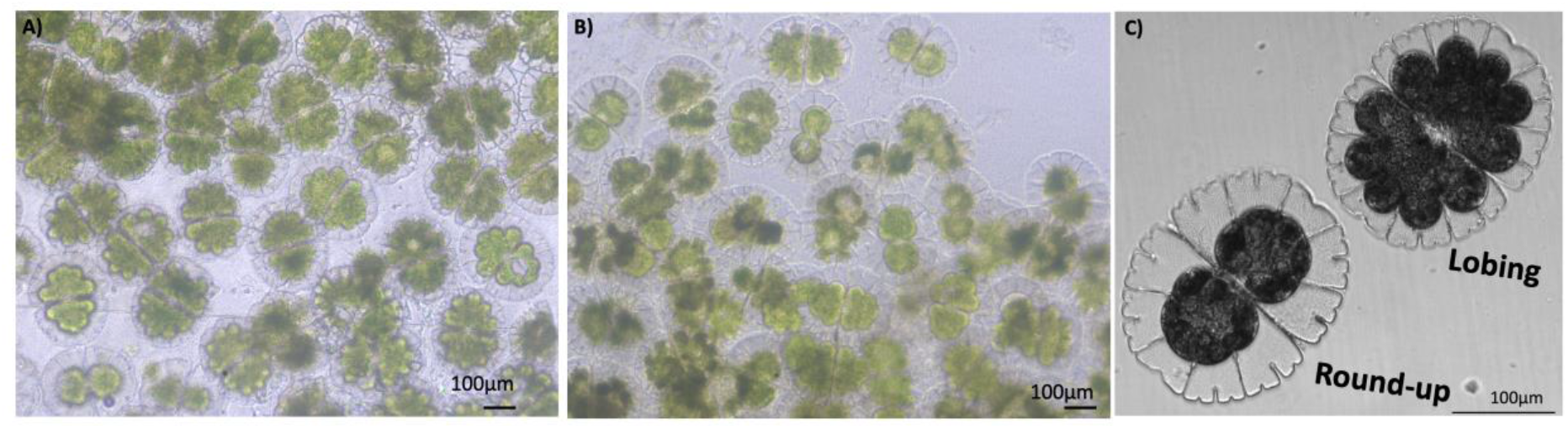
A) Cells in lobing stage in the induction medium with 5% mannitol, image taken after 5 hours. B) Cells in round up stage and few in lobing stage after 24 hours. C) Closeup view of lobing and round up stage. At the isthmus the protoplast rounds up, gets locked and remain at this stage.

Similar situation was noticed when the cells with induction medium were incubated at 37 **°**C. Overall, it appears that the use of mannitol and cellulysin as described in Berliner and Wenc protocol induces protoplast in *M.denticulata*, but they gets arrested at later stage within the semicell without getting released.

### Temperature ramping release intact protoplast from *M. denticulate* arrested in round up stage

After overnight incubation at 22 **°**C, the cells were shifted to 37 **°**C for two hours, left undisturbed and brought back to 22 **°**C for examination. *M. denticulata*, cells at 4 and 5% mannitol were found to be at different stages of actively releasing protoplast into the induction medium (Figure 3A). Thus, this shift in temperature for a short period seems to aid the cells to advance from the arrested round-up stage to protoplast emergence and release stage. As noted in other desmids, at the isthmus a membranous bubble appears and the green content of each semicell slowly moves into this bubble (Figure 3B-J). Bubble at isthmus occur from cells both in lobed and round up stages. Subsequently, when all the cell content got transferred into the bubble, this bubble becomes the protoplast and remain attached to one of the semicell, while the other gets detached. Eventually, both empty semicells get detached and round dark green intact protoplasts were released in the induction medium (Figure 3K). This release of protoplast did not happen in other concentrations of mannitol in this study, except at 4 and 5% mannitol treatment and this was confirmed by 5 repeated protoplast induction experiments. Not all the cells yielded protoplast in this temperature ramping strategy. The full protoplasts are roughly around 80-100 μm size and they remain round, unbroken in the induction medium for atleast 1 day at room temperature.

**Figure 3:**
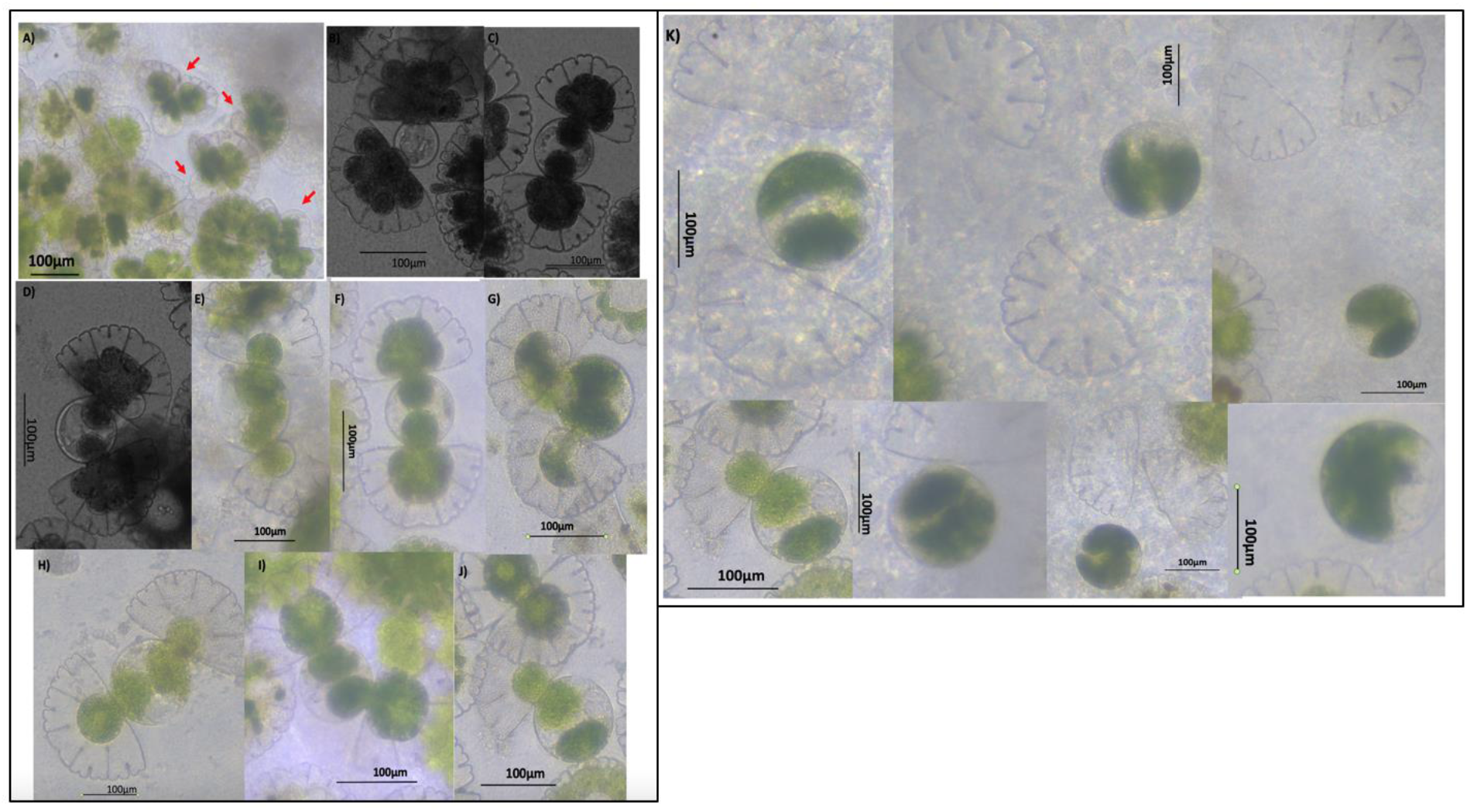
A) Cells after temperature ramp, actively engaged in bubble formation and growth are marked in red arrows. B-J) represents stages of bubble formation, bubble growth, entry of green content from semicells, filling up of the bubble to form future protoplast. Note that, as bubble grows in between them, the semicell move in backward direction. K) Snapshots of full protoplast released or getting released into the induction medium and the empty semicells.

In few cells, one of the semicell proceeds to round-up stage but the other remain in lobing stage (Figure 4A). In some cells, after retraction of membrane from the cell wall, during lobing, the lobes get fragmented within the semicells. In some other cells, not all the full content of semicells get transferred to the bubble at isthmus (Figure 4B). In such situations, the cells release protoplasts as small spherical units or subprotoplasts less than 40-60 μm size, that could be anucleate. Subprotoplasts were also found to be stable in the induction medium after release (Figure 4C).

**Figure 4:**
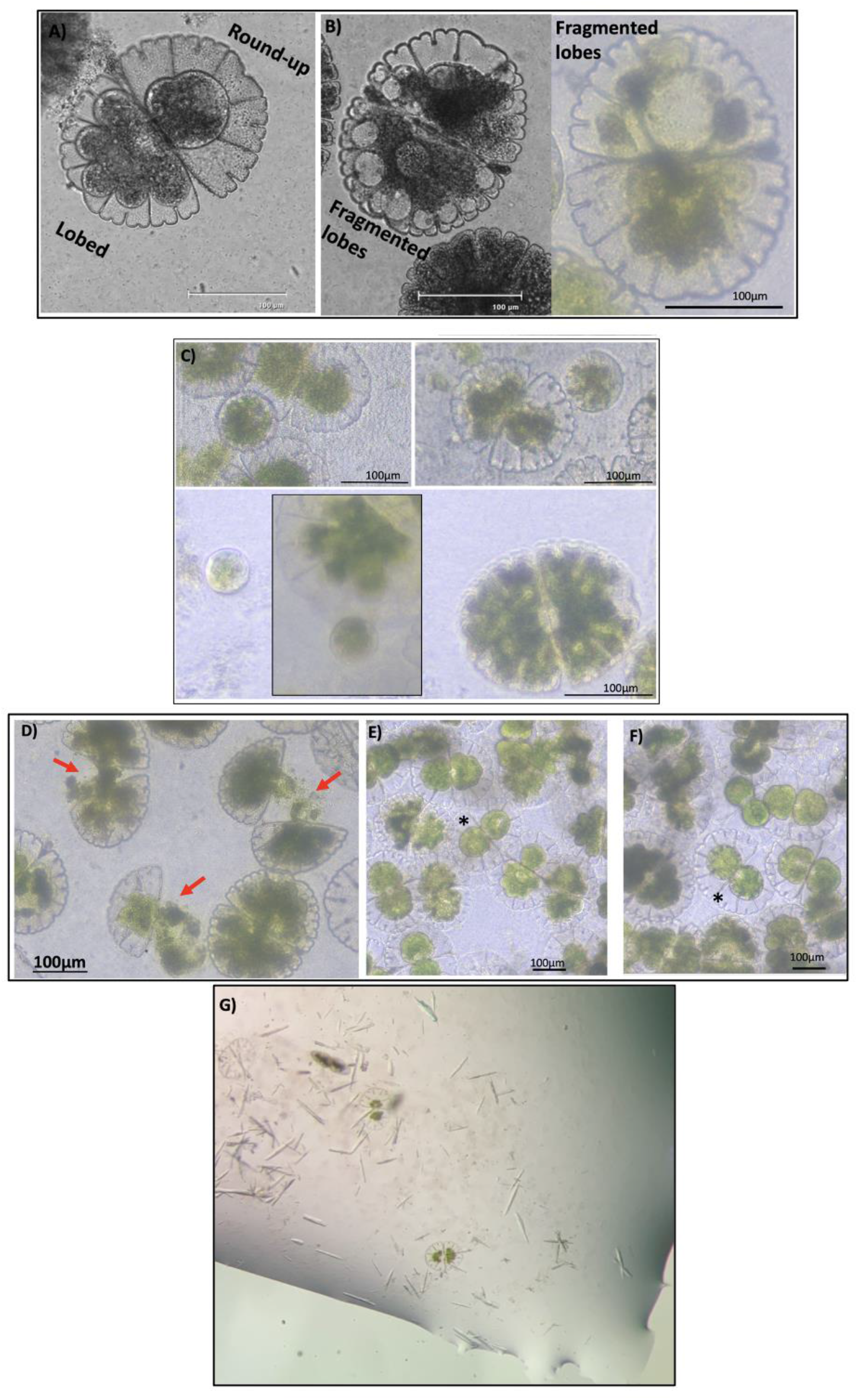
A) A cell with each of its semicell in different stages of progression to protoplast production. B) The two cells represent fragmentation of lobes inside each semicell as small circular vesicles during plasmolysis. C) Representative mini or subprotoplasts released from the above two situations. D). In the absence of calcium, many cells are found to have broken bubbles, highlighted by red arrows. E-F) A situation where a cell at its round-up stage surrounded by neighbours, locking it in a crowded environment. G) Needle like crystals found in the induction medium after overnight incubation with alga under 6% cellulysin.

Although protoplast could be produced without inclusion of calcium, it was noted that in the absence of 4mM CaCl2 there is high frequency of aborted protoplast production (Figure 4D). In the absence of calcium, the bubble that pops out of many cells at isthmus, breaks-off in the middle of transfer of materials from semicells and then, there is no further progress in protoplast formation from that cell.

### Crystal growth in the protoplast induction reaction

In a setup to study the effect of increasing cellulysin concentration on protoplast induction as the cells arrested in round-up stage, an induction assay at 22 **°**C with 6% cellulysin, 5% mannitol mixed with algae in distilled water was conducted. On overnight incubation, no protoplast was produced in this condition, but a number of needle like rectangular crystals appeared in the induction medium (Figure 4G). These crystals could be reproduced. Cellulysin powder is a cocktail of crude cellulase enzymes purchased from Sigma. The crystals are unlikely mannitol as there is no crystal growth even at 8-9% mannitol used in this study. In the protoplast induction reaction here, a concentrated protein sample (6% cellulysin (60mg/ml conc), a precipitant (mannitol), a salt (CaCl_2_) and the cellulosic algae are mixed and left undisturbed. This setup is a ‘batch method’ of protein crystallization, a past method to grow protein crystals, when the target protein is available in excess. It is difficult to say anything definite about the chemical nature of these crystals using light microscopy, unless it is analyzed by X-rays or electrons. These crystals were picked in nylon loop and frozen for a later full X-ray structure determination. It could be possible that at high concentration, the cellulase enzyme of fungus *Trichoderma viride* (source of cellulysin) got crystallized here in presence of its cellulosic substrate and in that case, structure determination from these crystals may represent a cellulase enzyme conformation in substrate or product fragment bound form.

## Discussion

The protocol described here expands the previously available method of protoplast production from desmids to suit *M.denticulata*. Here is a notable finding that a temperature ramp is required to release the protoplast from *M.denticulata* from its arrested round-up stage. With this, it is now becoming possible to produce intact protoplast from this alga in a simple two-step process. Obtaining good protoplasts from this model organism of cell wall research is essential for obtaining pure plasma membrane for high-resolution structural studies. Isolating plasma membrane, spherical and flat vesicles from this protoplast for cryo-electron microscopy has potential to open up structural molecular biology of cell wall biogenesis in *M.denticulata*, an algae evolutionarily close to land plants.

In further investigations, it would be interesting to probe along the following lines. Not all the cells yielded protoplast when subjected to this temperature ramping strategy at 4-5% mannitol concentration, but a countable number of protoplast could be produced. The underlying reasons for this are less clear. Since the algal culture used here is unsynchronized, only cells at a particular physiological state might be opt for complete protoplast release. Using synchronized culture for protoplast induction may clarify this. It is possible that the composition of isthmus surface varies between the cells, such that the cellulose there is accessible to cellulysin only in few cells and remain covered by other non-cellulosic polysaccharide in other cells. Inclusion of other wall material degrading enzymes in combination with cellulysin will be helpful in that situation.

After rounding up of protoplast, the plasma membrane pops out a transparent bubble from the isthmus, and this bubble grows in size as the content of each semicell enters it to form a protoplast. In this time, as the bubble grows the distance between two semicells increases and these semicells move away from each other backwards from the isthmus as their green content moves front and enters into the bubble. The flat disc shaped algae mostly settle at the bottom of the culture plate and at high concentration, each cell is in physical contact with its neighbour (Figure 4E-F). Cells in this crowded/tightly packed environment may experience difficulty for the necessary backward movement of semicells during bubble growth, influencing its emergence or growth, allowing only few cells to induce protoplast. On occasional stirring and on attempting protoplast induction in low number of cells, it was noticed that the crowding is not a factor controlling the number of protoplast release.

The sequence of events that proceeds from protoplast induction to release is apparent (Figure 5). But the mechanism underlying the protoplast release in this treatment of 4-5% mannitol and 2% cellulysin with a temperature ramp is less clear to the author. The following things have to be considered while framing a mechanism of protoplast release here in *M. denticulata*. Protoplast could be released only when the mannitol concentration is between 4 to 5% after the temperature change, although a range of mannitol concentration induces protoplast retraction from walls, lobing and rounding up of protoplast. In a limited range of temperature, an enzyme catalyzed reaction rate increases with temperature, but incubating the cells from the beginning at 37 **°**C does not result in protoplasts. This indicates that enhanced wall hydrolysis reactions by cellulysin at 37 **°**C is not the reason for the formation of protoplasts. Temperature ramping is a procedure in inducing large arrays of two-dimensional membrane protein crystals for electron crystallography (Kühlbrandt, 1992; Abeyrathne *et al.*, 2012). In this procedure, the protein crystallization set up is moved between room temperature and 37 **°**C once or repeatedly over a period of many hours, if the protein under study is stable. Mannitol operates by taking out water from the protoplast, making its water potential more negative with increasing concentration of mannitol, thereby altering its physical property. The strength of hydrophobic interaction increases with temperature (Tanford, 1980) and this influences the fluidity of the plasma membrane. It could be that, this temperature dependent membrane property coupled to the water potential of protoplast of certain cell stages at 4 to 5% mannitol is optimally suited for the successful emergence and release of protoplast in a temperature ramp through the cellulysin induced fractures/weak points at the isthmus of *M.denticulata* in this protoplast induction study.

**Figure 5:**
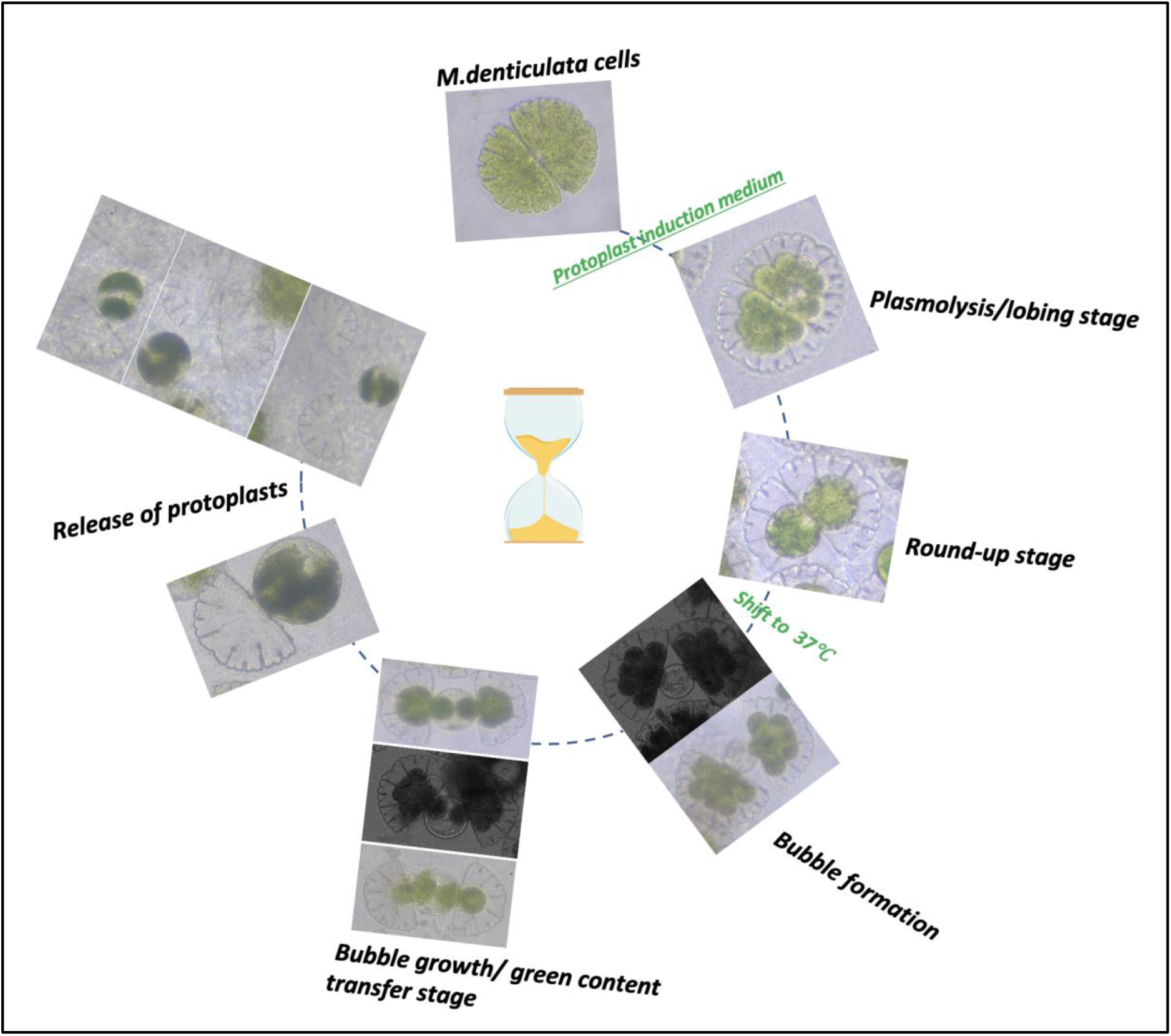
Sequence of events in the protoplast production in *M.denticulata*

## Acknowledgement

I thank Prof. Ursula Lütz-Meindl, University of Salzburg for generously agreeing to familiarize me with growth and development of this alga (*M. denticulate*) through a short stay in her lab, but unfortunately passed away before the visit (Holzinger, 2020). I am grateful to her former graduate students Prof. Andreas Holzinger, University of Innsbruck, Dr.Phillip Steiner and her lab manager Mag.Ancuela Andosch for sending me this algae and remotely guiding on its cultivation. I thank Dr.Timo Saarinen, University of Helsinki for granting me access to their plant growth facility. I am very much grateful to Prof. Pekka Lappalainen for his support to conduct this study in his laboratory. I thank Dr.Indu Santhanagopalan, University of Cambridge for her suggestions and critical comments in this.

## Supplementary material

### Composition of Desmidiacean medium

(provided by Ancuela Andosch, University of Salzburg, Austria)

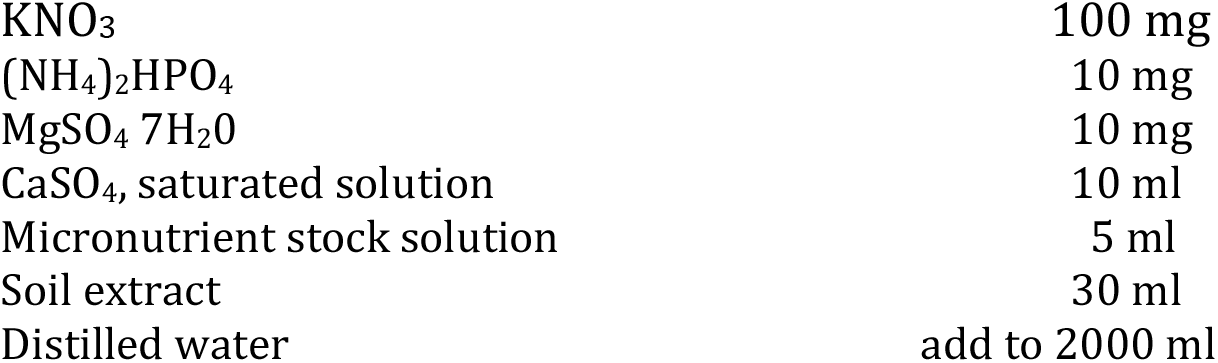

|  |  |
| --- | --- |
| KNO<sub>3</sub> | 100 mg |
| (NH<sub>4</sub>)<sub>2</sub>HPO<sub>4</sub> | 10 mg |
| MgSO<sub>4</sub> 7H<sub>2</sub>O | 10 mg |
| CaSO<sub>4</sub>, saturated solution | 10 ml |
| Micronutrient stock solution | 5 ml |
| Soil extract | 30 ml |
| Distilled water | add to 2000 ml |

In each 30ml culture media of a 100ml flask, a drop of 0.001% sterile Vitamin B12 solution is added and inoculated with algae. A light:dark regime of 14:10 hours at 20°C, with light intensity between 100-150μmol photons.m^-2^.s^-1^ is preferred. Subculturing at every 3-4 weeks is recommended.

### Micronutrient stock solution

Micronutrient stock solution:
|  | <u>Stock solution</u> | <u>Handling Solution</u> |
| --- | --- | --- |
| ZnSO<sub>4</sub> 7 H<sub>2</sub>O | 0,1% | 1 ml |
| MnSO<sub>4</sub> 4 H<sub>2</sub>O | 0,1% | 2 ml |
| H<sub>3</sub>BO<sub>3</sub> | 0,2% | 5 ml |
| Co(NO<sub>3</sub>)<sub>2</sub> 6 H<sub>2</sub>O | 0,02% | 5 ml |
| Na<sub>2</sub>MoO<sub>4</sub> 2 H<sub>2</sub>O | 0,02% | 5 ml |
| CuSO<sub>4</sub> 5 H<sub>2</sub>O | 0,0005% | 1 ml |
| Aqua dest. |  | 981 ml |
| Fe SO<sub>4</sub> 7 H<sub>2</sub>O |  | 0,7 g |
| EDTA (Titriplex III) |  | 0,8 g |

The components need to be prepared in as below two separated solutions and autoclaved. After cooling both solutions need to be mixed under sterile conditions.

**Solution 1**: 881 ml distilled water + 19 ml Micronutrient stock solution (without Fe) + 0,4 g EDTA

**Solution 2**: 100 ml distilled+ 0,7 g FeSO_4_ 7 H_2_0 + 0,4 G EDTA

